# Ultrafast endocytosis in mouse cortical inhibitory synapses

**DOI:** 10.1101/2025.06.06.658279

**Authors:** Chelsy R. Eddings, Shigeki Watanabe

## Abstract

Neural circuitry depends on excitatory and inhibitory regulation of activity to yield functional outputs. However, examinations of synaptic vesicle recycling have focused heavily on excitatory synapses, leaving many questions unanswered for inhibitory synapse dynamics. Here we show that both excitatory and inhibitory cortical synapses contain depots of a protein essential for ultrafast endocytosis, Dynamin 1xA, at a region immediately next to the active zone where ultrafast endocytosis takes place. Using zap-and-freeze time-resolved electron microscopy in mouse acute cortical slices, we observe uncoated pits reminiscent of ultrafast endocytic intermediates appearing post-stimulus at putative inhibitory synapses. These findings suggest that excitatory and inhibitory synapses may perform similar modes of endocytosis.

## Introduction

Inhibitory synapses are essential for brain function. Disruption of cortical excitatory-inhibitory synapse balance can result in developmental, psychiatric, and neurodegenerative disorders. For example, postmortem autism patient brains contain increased excitatory synapses in upper cortical layers, alongside decreased inhibitory synapses across all cortical layers^1^. Postmortem schizophrenia dorsolateral prefrontal cortex displays several complex alterations in inhibitory interneurons and deficits in presynaptic γ-aminobutyric acid (GABA) synthesizing enzymes like GAD67^2^. Alzheimer’s postmortem parietal cortex exhibits decreased gephyrin-positive inhibitory synapses in layer 2, likely contributing to hyperactivity and intellectual decline^3^. Therefore, it is essential to have a thorough understanding of inhibitory synapse dynamics.

Existing studies of synaptic vesicle dynamics highlight potential differences between excitatory and inhibitory synapses. Within the hippocampus, minutes after chemical or ischemia-like stimulation, both excitatory and inhibitory synapses exhibit vesicle depletion as well as increased clathrin-coated endocytic pits and vesicles^4^. Although the rates of exocytosis are similar, the kinetics of endocytosis are ∼80% slower in inhibitory synapses, when measured by pHluorin at 26 °C^5^. This difference is more subtle but evident even when measured at 35 °C^5^. Differences between excitatory and inhibitory subtypes also exist at the level of individual synaptic vesicles. GABAergic VGAT-labeled vesicles travel less distance in a more direct route to fusion sites in the presynaptic active zone and undergo exocytosis earlier than Synaptotagmin-1 labeled vesicles^6^. GABAergic vesicles also have higher resting luminal pH and display an overacidification following endocytosis—requiring later alkalization to restore internal pH after the overshoot^7^. Additionally, GABAergic synaptic vesicles show faster initial proton efflux, but slower vesicle reacidification compared to glutamatergic vesicles^8^. These differences likely reflect the need for inhibitory synapses to rapidly produce high-fidelity signaling within complex neural circuits.

Nonetheless, the mechanisms of synaptic vesicle endocytosis in inhibitory synapses have not been well defined. To date, five different modes of endocytosis are described in neurons: clathrin-mediated endocytosis, kiss-and-run, ultrafast endocytosis, fast compensatory endocytosis, and activity-dependent bulk endocytosis. These modes are separated by their reliance on clathrin machinery, location, and timing^9^. Clathrin-mediated endocytosis occurs outside of the active zone, within 10-30 seconds after stimulation^10,11^. Kiss-and-run recycles vesicles via the reversal of a fusion pore directly at the active zone and completes potentially as fast as 10 milliseconds but within 1-2 seconds^12–18^. Ultrafast endocytosis occurs immediately next to the active zone, internalizing excess membrane as quickly as 50 milliseconds up to ∼1 second^19,20^. Compensatory endocytosis has kinetics similar to ultrafast endocytosis, and may share similar mechanisms, but its location is unknown^21,22^. Finally, bulk endocytosis, triggered by intense stimulation, occurs outside of the active zone within seconds to minutes^17,18^. Over the last decade, we have demonstrated the existence of ultrafast endocytosis in neurons from *Caenorhabditis elegans*^19^, mice^20^, and humans^23^, but ambiguity remains for the modes of endocytosis present in inhibitory synapses.

Here we leverage zap-and-freeze electron microscopy (EM) to probe the membrane dynamics of mouse cortical synapses. Zap-and-freeze utilizes electric field stimulation of neurons and high-pressure freezing to yield millisecond and nanometer observations of membrane trafficking events^24^. We investigate inhibitory synapses on a rapid timescale to determine if our previous findings, suggesting the presence of ultrafast endocytosis in the mouse cortex^23^, are generalizable across excitatory and inhibitory synapse subtypes. Our results in acute cortical slices indicate the possible presence of ultrafast endocytosis in inhibitory synapses after a single stimulus. Furthermore, a splice variant of Dynamin 1, Dynamin 1xA (Dyn1xA)^25,26^, localizes immediately next to the active zone where uncoated endocytic pits form. These data suggest that ultrafast endocytosis also recycles synaptic vesicles in inhibitory synapses.

## Materials and Methods

### Mice

All procedures involving mice were approved by the Johns Hopkins Animal Care and Use Committee and followed the guidelines of the National Institutes of Health. Both males and females were used in this study. All animals were kept on a 12 hour light/dark cycle and provided access to unlimited food and water. Wild-type mice used were obtained from Charles River (C57BL/6) and Taconic (B6NTAC).

### Primary neuron cultures

Primary cortical cultures were prepared from embryonic day 18 (E18) or newborn (P0) pups of unidentified genders. Pups were decapitated, with their brains transferred to ice cold dissection medium (1x HBSS, 1 mM sodium pyruvate, 10 mM HEPES, 30 mM glucose, and 1% penicillin-streptomycin). Cortices were dissected under a binocular microscope and digested with papain (0.5 mg/ml, Worthington Biochemical, cat#: LS003119) and DNase (0.01%, Millipore Sigma, cat#: DN25) in dissection medium for 20 min at 37 °C. Cells were then further dissociated by trituration using fire-polished Pasteur pipettes.

For immunocytochemistry, neurons were seeded on coverslips (18 mm, #1.5H thickness, Neuvitro Corporation, cat#: GG-18-15H) that had been previously coated overnight with poly-L-lysine (1 mg/mL, Sigma-Aldrich, cat#: P2636), at a density of 125 × 10^3^ cells per well of a 12-well plate (Corning).

For high-pressure freezing, neurons were cultured on 6-mm sapphire disks (Technotrade International, cat#: 616-100) coated with poly-D-lysine (1 mg/ml, Sigma-Aldrich, cat#: P6407) and rat tail collagen I (0.6 mg/ml, ThermoFisher, cat#: A1048301), and pre-seeded with an astrocyte feeder layer. Astrocytes were harvested at least 3 weeks beforehand from E18/P0 mouse cortices with treatment of trypsin (0.05%, ThermoFisher, cat#: 25300054) for 20 min at 37 °C, followed by trituration. After 2 weeks, astrocytes were plated onto sapphire disks at a density of 50 × 10^3^ cells per well. After 1 week, astrocytes were incubated with 5-Fluoro-2′-deoxyuridine (81 μM; Sigma-Aldrich, cat#: F0503) and uridine (204 μM; Sigma-Aldrich, cat#: U3003) for at least 2 hours to stop mitosis, and then medium was switched to Neurobasal-A (Gibco) supplemented with 2 mM GlutaMax, 2% B27 and 0.2% penicillin-streptomycin prior to addition of neurons. Neurons were seeded at a density of 75 × 10^3^ cells per well of a 12-well plate (Corning).

### Acute brain slice preparation

Acute mouse cortical slices were cut to 100 µm on a vibratome (Leica VT1200S) in room temperature NMDG aCSF (in mM: 92 NMDG, 2.5 KCl, 1.25 NaH_2_PO4, 30 NaHCO_3_, 20HEPES, 25 glucose, 2 thiourea, 5 Na-ascorbate, 3 Na-pyruvate, 10 NAC, 0.5 CaCl_2_·2H_2_O, and 10 MgSO_4_·7H_2_O). Brain slices were then transferred into a custom-built recovery chamber filled with continuously carbogenated NMDG solution at 37 °C and allowed to undergo an initial 12 min recovery. After 12 min, slices were moved into a new recovery chamber filled with continuously carbogenated HEPES holding aCSF (in mM: 92 NaCl, 2.5 KCl, 1.25 NaH_2_PO_4_, 30 NaHCO_3_, 20 HEPES, 25 glucose, 2 thiourea, 5 Na-ascorbate, 3 Na-pyruvate, 2 CaCl_2_·2H_2_O, and 2 MgSO_4_·7H_2_O) and allowed to undergo a second recovery for at least 4 hr at 37 °C. Slices were recovered to ensure electrical viability and responsiveness to later electric field stimulation—also to repair cut site damage to maximize the amount of maintained tissue morphology while imaging. All aCSF solutions were titrated to 7.3-7.4 pH, left at room temperature to avoid potential tissue shrinkage-expansion cycles that can affect tissue ultrastructure, and bubbled with carbogen gas (95% O_2_, 5% CO_2_) before use. Note that we did not perform the Na+ spike-in procedure associated with the original NMDG recovery method^27^ to prevent possible tissue overexcitation.

### Zap-and-freeze experiments

For zap-and-freeze of primary mouse neurons, DIV 14 cortical cultures were used. Neurons on sapphire disks were placed in physiological saline solution containing 140 mM NaCl, 2.4 mM KCl, 10 mM HEPES, 10 mM Glucose (pH 7.3), 300 mOsm, 4 mM CaCl_2_, and 1 mM MgCl_2_. Additionally, NBQX (3 mM, Tocris, cat#: 1044) and Bicuculline (3 mM, Tocris, cat#: 0109) were added to suppress recurrent network activity following electrical stimulation of neurons. Solutions and equipment were kept at 37 ºC before freezing.

For ferritin-loading experiments, cationized ferritin from horse spleen (Sigma-Aldrich, cat#: F7879) was added to the saline solution at 0.25 mg/ml. Calcium concentrations were reduced to 1 mM to suppress spontaneous activity during loading. Cells were incubated in ferritin solution for 5 min at 37 ºC on a shaker. After incubation, the cells were immersed in fresh saline solution containing 4 mM Ca^2+^ for less than 2 min before being assembled into the freezing chamber of the Leica EM ICE high-pressure freezer.

For zap-and-freeze of acute mouse brain slices, tissue was always tested the same day as collection. Slices were transported from recovery chambers to the high-pressure freezer in a petri-dish filled with room temperature aCSF (in mM: 125 NaCl, 2.5 KCl, 1.25 NaH_2_PO_4_, 25 NaHCO_3_, 10 glucose, 2 CaCl_2_·2H_2_O, and 2 MgSO_4_·7H_2_O; 7.3-7.4 pH). Slices were then trimmed into smaller pieces ∼6 mm by hand using a razor blade. After trimming, slices were placed into freezing medium containing pre-warmed aCSF, supplemented with NBQX (3 mM), Bicuculline (3 mM), and 15% polyvinylpyrrolidone (PVP) as a cryoprotectant—slices were kept in this solution for less than 2 min as the freezing apparatus was assembled. 15% PVP was chosen and assembled in a specimen sandwich based on a method previously published for optogenetic stimulation of mouse brain slices using the high-pressure freezer^28^. The table and sample chamber of the high-pressure freezer were kept at 37 ºC to ensure the physiological temperature of slices were maintained during experiments^29^. Unstimulated controls for each experiment were always frozen on the same day and originated from the same mouse or patient. The device was set such that samples were frozen at 0.1 or 1 sec after the stimulus initiation. Cortical columns were always visually aligned perpendicular to the direction of the electric field to maintain consistency across samples (this ensured cortical layers were in parallel with the electric field^23^).

### Freeze substitution

Frozen samples were transferred under liquid nitrogen to an automated freeze substitution unit held at -90 °C (EM AFS2, Leica Microsystems). Using chilled tweezers, samples were moved into acetone to help disassemble the freezing apparatus. Sapphire disks with brain slices were then quickly moved into sample carriers containing 1% glutaraldehyde (GA) and 0.1% tannic acid (TA) in anhydrous acetone. The freeze substitution program was as follows: -90 °C for at least 36 hrs with samples in 1% GA, 0.1% TA, samples were then washed five times with pre-chilled acetone (30 min each), after washing the fixative solution was replaced with pre-chilled 2% OsO_4_ in acetone and the program was allowed to continue; -90 °C for 11 hrs with samples in 2% OsO_4_, +5°C per hour to -20 °C, -20 °C for 12 hrs, +10 °C per hour to +4 °C, hold at +4 °C.

### Sample preparation for electron microscopy

After freeze substitution, samples removed from the AFS were washed six times with acetone (10 min each) and incubated with increasing levels of plastic (100% epon-araldite diluted with acetone: 30% for 2 hrs, 70% for 3 hrs, and 90% overnight at +4 °C). After plastic infiltration, samples were embedded in 100% epon-araldite resin (Araldite 4.4 g, Eponate 12 Resin 6.2 g, Dodecenyl Succinic Anhydride (DDSA) 12.2 g, and Benzyldimethylamine (BDMA) 0.8 ml) and cured for 48 hrs in a 60 °C oven. Serial 40-nm sections were then cut using an ultramicrotome (EM UC7, Leica microsystems) and collected onto 0.7% pioloform-coated single-slot copper grids. Sections were stained with 2.5% uranyl acetate in a 50-50 methanol-water solution.

### Transmission electron microscopy

Images were acquired on a Hitachi 7600 transmission electron microscope at 80 kV and 80,000x magnification, with a AMT XR80 high-resolution (16-bit) 8-megapixel CCD camera, using AMT Image capture engine V602 (AMTV602) software. Samples were blinded and given random names before imaging. At least 98-100 images of random cortical synapses were obtained per experimental timepoint. For specific inhibitory synapse analysis, at least 47 images were obtained per experimental timepoint. Inhibitory synapses were identified in electron micrographs as presenting a non-prominent postsynaptic density, with synaptic connections onto either cell bodies or being on dendritic shafts^30–33^.

### Electron micrograph image analysis

Electron micrographs were manually analyzed using the published SynapsEM protocol^34^. Images from one experimental replicate were pooled into a single folder, randomized, and blinded using Matlab scripts. Synapses not containing a prominent postsynaptic density or those with poor morphology were manually excluded from analysis after this blinded randomization. Using custom Fiji macros, membrane and organelle features were annotated and exported as text files. Those text files were again imported into Matlab where the number and locations of the annotated features were calculated. For pit distribution from the active zone, the distance from the nearest edge of pits to the active zone was calculated. To minimize bias and error all annotated, randomized images were thoroughly checked and edited by at least one other member of the lab. Representative electron micrographs were adjusted in brightness and contrast to different degrees, rotated and cropped in Adobe Photoshop (v21.2.1) or Illustrator (v24.2.3). All Fiji macros and Matlab scripts are publicly available at https://github.com/shigekiwatanabe/SynapsEM.

### Immunocytochemistry

On DIV 14 cortical neurons cultured on coverslips (18 mm, #1.5H thickness, Neuvitro Corporation, cat#: GG-18-15H) were fixed with 4% paraformaldehyde (PFA; Electron Microscopy Sciences, cat#: 15714) in PBS for 20 min at room temperature. Cells were then washed with PBS, permeabilized with 0.2% Triton X-100 in PBS for 8 min, washed again with PBS, and incubated with 10% Normal Goat Serum (NGS, Jackson ImmunoResearch, cat#: 005-000-121) in PBS for 1 hr of blocking. The cells were then incubated overnight at +4 °C in PBS containing 10% NGS and primary antibodies. Primary antibodies included: Dynamin1xA (rabbit, 1:500), Bassoon (mouse, 1:500), FluoTag-X2 anti-PSD95 (1:50), and Gephyrin (guinea pig, 1:100). The next day, cells were washed with PBS and incubated with secondary antibodies diluted in 10% NGS in PBS for 1 hr—for excitatory synapses: anti-rabbit Alexa 594 (1:500) and STAR 460L goat anti-mouse IgG (1:150); for inhibitory synapses: anti-rabbit Atto 647 (1:200), anti-mouse Alexa 594 (1:500), and STAR 460L goat anti-guinea pig IgG (1:150). PBS washed coverslips were finally dipped in Milli-Q water using forceps, dried by tapping on a filter paper, and mounted onto plain microscope slides with ProLong Diamond Antifade Mountant (Invitrogen, cat#: P36970). Samples were allowed to dry at least overnight before STED imaging.

### Immunohistochemistry

For immunohistochemistry, adult mice were anesthetized with isoflurane, transcardially perfused with 4% paraformaldehyde (PFA) and decapitated, with their whole brain dissected into PBS. Whole brains were embedded in O.C.T. compound (Tissue-Tek), frozen on dry ice, and sectioned in a coronal orientation at 40 µm on a cryostat (Leica CM 3050S). Staining was performed on free-floating sections according to the protocol described in Kruzich, E, et al^35^. Slices were permeabilized and blocked with 10% Normal Goat Serum (NGS) and 1% Triton X-100 in PBS for 3 hrs at room temperature. Primary antibodies diluted in 10% NGS and 0.025% Triton X-100 in PBS were added to slices for 48 hrs at +4 °C. Primary antibodies for inhibitory synapses included: Bassoon (mouse, 1:100), Dynamin1xA (rabbit, 1:100), and Gephyrin (chicken, 1:100). After removal of primary antibodies, slices were washed four times with 0.025% Triton X-100 in PBS, 15 min each wash. Secondary antibodies were diluted in PBS and incubated with slices for 48 hrs at +4 °C. Secondary antibodies included: STAR RED goat anti-mouse IgG (1:200), STAR ORANGE goat anti-rabbit IgG (1:100), and STAR 460L goat anti-chicken IgY (1:50). Slices were washed four times with PBS, then washed once with Milli-Q water before being mounted onto plain microscope slides (Globe Scientific) using a paintbrush. Tissue was allowed to dry until transparent under foil. ProLong Diamond Antifade Mountant was added directly onto slices, with a coverslip placed on top. Samples were left to dry at least overnight before STED imaging.

### Stimulated emission depletion (STED) imaging

2D STED images were acquired on an Aberrior FACILITY line microscope using a 60x oil objective lens (NA = 1.42). The excitation wavelengths were set as: 640 nm, 561 nm, and 485 nm for imaging Atto-643/Atto-647/STAR-RED, Alexa-594/STAR-ORANGE, and STAR460L labeled target respectively. Imaging was performed at 20 nm pixel size, 1.0 AU pinhole for cultures and 0.61 AU for slices, dwell time 5 µs. The STED beam was set at 775 nm with power of 10-15% used.

### STED image deconvolution, segmentation and analysis

STED images were exported in .obf format using LiGHTBOX software (Abberior). The remaining image processing was performed using a custom made MATLAB code package (Imoto et al. 2024^26^). Images were extracted from .obf files and converted to .tif format as unsigned 16-bit integers. The extracted images were normalized based on the minimum and maximum intensity values within each image and then blurred with a Gaussian filter with 1.2 pixel radius to reduce the Poisson noise. Subsequently, the images were deconvoluted twice using the two-step blinded deconvolution method. The initial point spread function (PSF) input was measured from the unspecific antibody signals of STAR 460L, Alexa 594, STAR ORANGE, Atto 643, or STAR RED in the STED images. The second PSF (enhanced PSF) input was chosen as the returned PSF from the initial run of blind deconvolution^36^. The enhanced PSF was used to deconvolute the STED images to be analyzed. Each time 10 iterations were performed. Series of the deconvoluted STED images were loaded to the segmentation script utilizing MIJ: Running ImageJ and Fiji within MATLAB (Sage 2017, MATLAB Central File Exchange). All presynaptic boutons in each deconvoluted images were selected within 45×45-pixel regions of interest (ROIs) based on the Bassoon and Dyn1xA signals. Top view and side view presynapses are sorted via script with supervision based on Bassoon shape. Distance distribution analysis was performed on top view images only, since Dyn1xA puncta can appear to be in the middle of synapses when seen in side view.

The boundary of active zone or Dyn1xA puncta was identified as the contour that represents half of the intensity of each local maxima in the Bassoon channel. The Dyn1xA puncta were picked by calculating pixels of local maxima. The distances between the Dyn1xA puncta and active zone boundary were automatically calculated correspondingly. For this distance measurement, first, MATLAB distance2curve function (John D’Errico 2024, MATLAB Central File Exchange) calculated the distance between the local maxima pixel and all the points on the contour of the active zone or Dyn1xA cluster boundary. Next, the minimum distance for each local maxima pixel was selected. Signals crossing the ROIs and the Bassoon signals outside of the stained neurons were excluded from the analysis. The MATLAB scripts are available from GitHub (https://github.com/imotolab-neuroem/STED_image_analysis_package_public_v1.4) or by request.

## Results

### Dyn1xA clusters in excitatory and inhibitory cortical presynapses

Our recent work in excitatory cortical synapses from mouse and human brain slices suggests that ultrafast endocytosis is retrieving synaptic vesicles following a single stimulus^23^.Towards our goal of determining if a similar mechanism occurs in inhibitory synapses, we first localized Dyn1xA, a protein critical for ultrafast endocytosis. Using our recently developed antibody, recognizing a unique C-terminal 20 amino acid region on Dyn1xA^23^, we examined endogenous protein localizations using stimulated emission depletion (STED) microscopy. Analysis of cultured mouse cortical synapses showed punctate Dyn1xA antibody signals in both excitatory PSD95-positive and inhibitory gephyrin-positive synapses (Figure 1A; see Figure S1A-C for example cultured neuron STED overview and additional side view synapses that were present but not analyzed). Dyn1xA puncta were found directly at the active zone boundary and periactive zone (−50 nm to +50 nm) (Figure 1B), with nearly 36% and 27% of puncta localized within this region in excitatory and inhibitory synapses, respectively (Figure 1C). There were no significant differences in the distance distributions of Dyn1xA puncta when comparing PSD95-positive and gephyrin-positive synapses (for Figure 1B p=0.9934; for Figure 1C p>0.9999, both using Kolmogorov-Smirnov tests). Therefore, Dyn1xA is present at the reported ‘endocytic zone’^25^ of both excitatory and inhibitory mouse cortical synapses.

**Figure 1.**
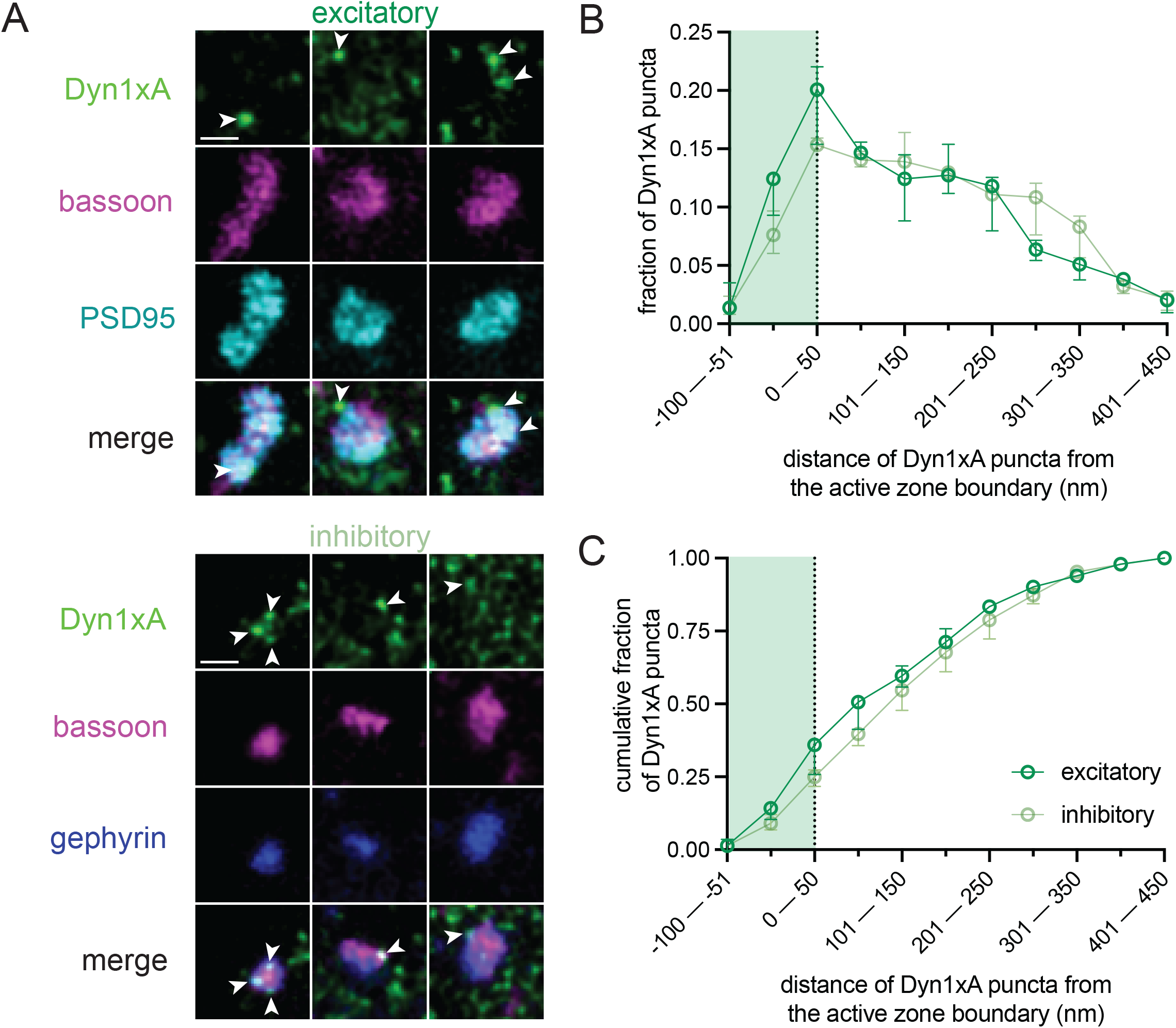
Dyn1xA clusters in excitatory and inhibitory cultured cortical presynapses. (A) Example STED images of endogenous Dyn1xA localizations (white arrowheads) in excitatory (PSD-95 expressing) and inhibitory (gephyrin-expressing) synapses of cultured mouse cortical neurons. Scale bars: 300 nm. More example STED images provided in Figure S1. (B) The distribution of Dyn1xA puncta relative to the active zone edge, defined by Bassoon, analyzed in top view synapse images. Shaded region indicates area inside the active zone Bassoon signal. The median and 95% confidence interval are shown for n=3 independent replicate cultures; 340 excitatory top view synapses and 297 inhibitory top view synapses were analyzed. (C)Cumulative plots of data presented in (B).

### Uncoated pits form 100 ms after stimulation in cultured cortical synapses

To confirm that ultrafast endocytosis occurs in mouse cortical neuron cultures, we performed zap-and-freeze EM. As in previous studies, cationized ferritin particles acted as fluid-phase markers of endocytosis^20,37^. After cells were incubated in ferritin containing media, a single 1 ms electric field stimulus was applied, and neurons frozen at 100 ms and 1 s. Unstimulated controls, also incubated in ferritin containing media, were always frozen on the same day from the same cultures. In electron micrographs, the active zone is defined as the presynaptic membrane region juxtaposed to the postsynaptic density. Excitatory and inhibitory synapses have been typically distinguished in EM by: the thickness of their postsynaptic densities (creating asymmetry or symmetry), the shape of their synaptic vesicles (circular or flatten), and their general connection location on neurons (dendritic spines versus cell bodies or dendritic shafts)^30,38–41^. However, cryo-preserved specimens do not show the robust vesicle flattening as seen in chemically fixed samples^42^. Also the distinction of postsynaptic density thickness is too subtle in cultured neurons. As such we could not verify the exact synapse types in our electron micrographs from cortical cultures. Nonetheless, cultured cortical presynapses showed uncoated pits forming 100 ms after a single electric field stimulus (Figure 2A,B; see Figure S2 for more EM images). These pits formed primarily 20-50 nm from the active zone (Figure 2C, median pit distances: 22.01 and 54.71 nm for culture 1 and 2 respectively). No clear clathrin-coated pits or Ω-figures were found in any of the tested timepoints. Ferritin-positive membrane structures like large vesicles (with diameter over 60 nm) and endosomes (with diameter 100 nm or more) increased in numbers one second after stimulation (Figure 2B). These results corroborate past findings in acute mouse brain slices^23^, that suggest indiscriminately tested cortical synapses can perform clathrin-independent endocytosis.

**Figure 2.**
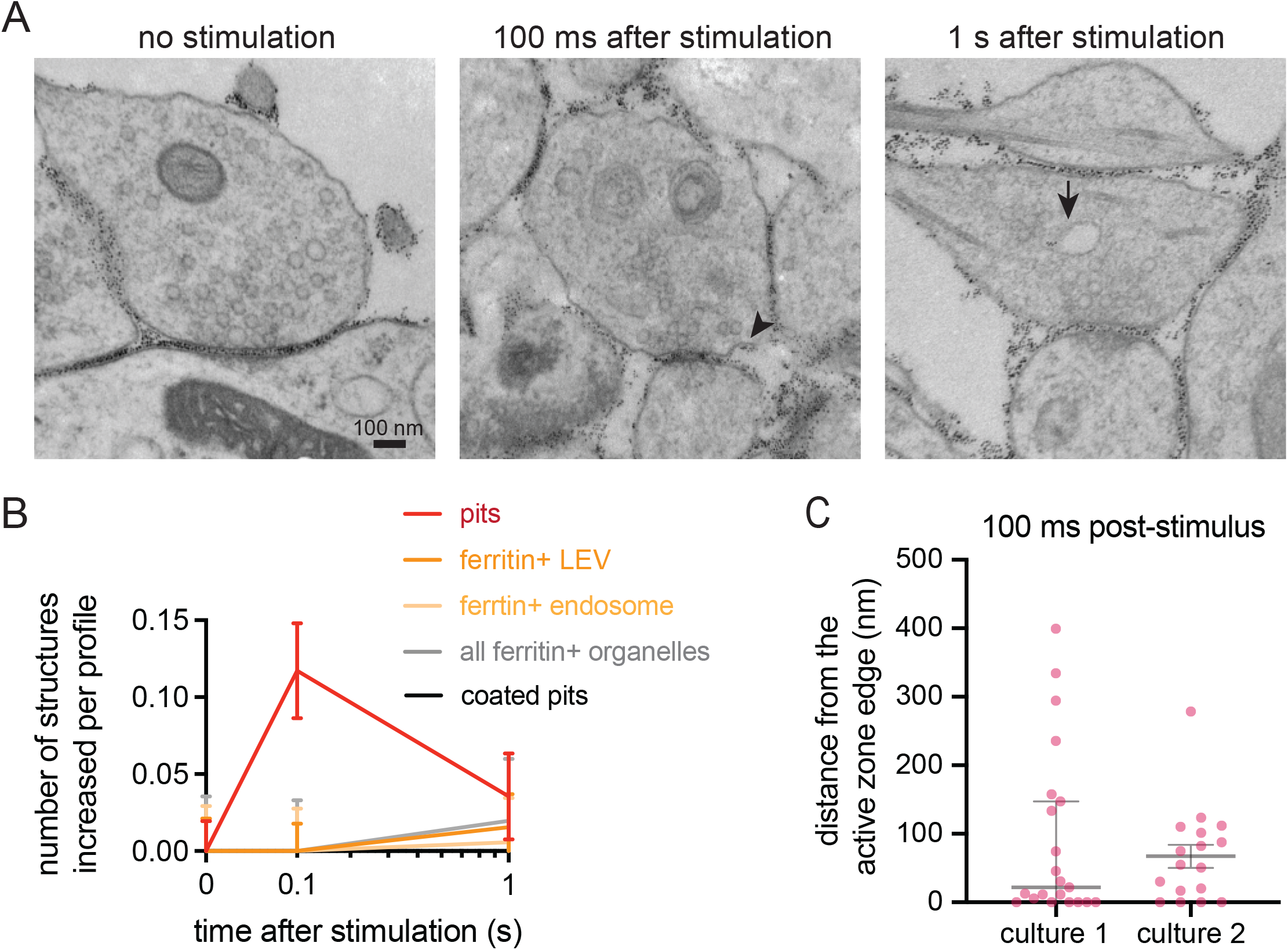
Uncoated pits form 100 ms after stimulation in cultured cortical synapses. (A) Example electron micrographs showing an uncoated endocytic pit (black arrowhead) and putative endosome (black arrow) at the indicated time points in DIV 14 cultured mouse cortical neurons. Here no distinction was made between excitatory or inhibitory synapses. Scale bar: 100 nm. More example TEM images are provided in Figure S2. (B) Plots showing the increase in number of each endocytic structure per synaptic profile after a single stimulus. Uncoated pits (pits), ferritin-positive large vesicles (ferritin+ LEV), ferritin-positive endosomes (ferritin+ endosome), ferritin-positive large vesicles and endosomes combined (all ferritin+ organelles) and clathrin-coated pits (coated pits) are indicated. Data are pooled from two independent cultures and presented as mean ± SEM. n values for each timepoint: no stim, n = 200 synapses; 100 ms, n = 203 synapses; 1 s, n = 199 synapses. (C) Plot showing the distance distribution of uncoated endocytic pits from the edge of an active zone 100 ms post-stimulus in cultured mouse cortical neurons. Data are presented as median ± 95% confidence interval. Each dot represents an uncoated pit.

### Dyn1xA clusters in inhibitory synapses within acute cortical slices

To assess inhibitory presynapses more selectively and within intact brain tissue, we next tested acute cortical slices. STED imaging from acute slices across three mice substantiated our primary neuron culture results—Dyn1xA puncta were found in gephyrin-positive synapses (Figure 3A; Figure S3 for example mouse slice STED overview and additional side view synapses that were present but not analyzed), located predominantly at the active zone boundary and periactive zone (−50 nm to +50 nm) (Figure 3B), with up to ∼41% of puncta localized in this region (Figure 3C). These data suggest that inhibitory synapses in an intact circuit context also contain a depot of Dyn1xA at the described endocytic zone.

**Figure 3.**
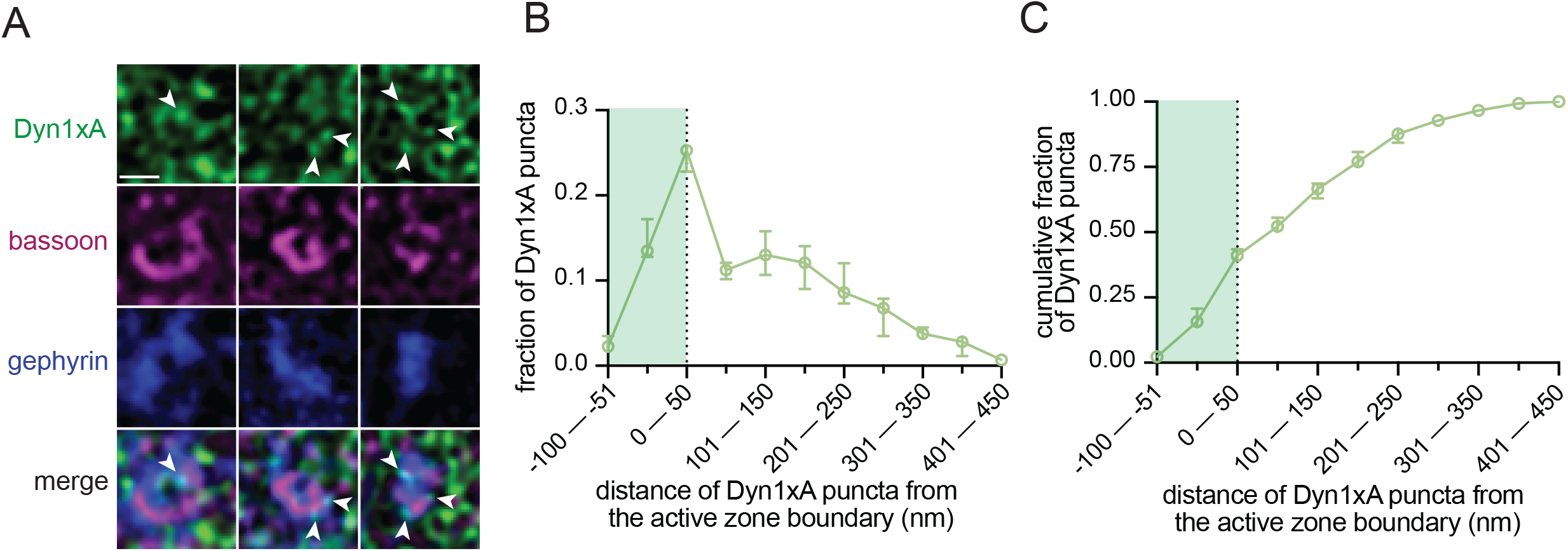
Dyn1xA clusters in inhibitory synapses within acute cortical slices. (A) Example STED images of endogenous Dyn1xA localizations (white arrowheads) in inhibitory (gephyrin-expressing) synapses of acute mouse cortical slices. Scale bar: 300 nm. More example STED images provided in Figure S3. (B) The distribution of Dyn1xA puncta relative to the active zone edge, defined by Bassoon, analyzed in top view synapse images. Shaded region indicates area inside the active zone Bassoon signal. The median and 95% confidence interval are shown for n=3 independent mice; 199 total inhibitory top view synapses were analyzed. (C) Cumulative plots of data presented in (B).

### Uncoated pits form 100 ms after stimulation in putative inhibitory synapses

Finally, we surveyed the ultrastructural dynamics of activated inhibitory synapses by repurposing zap-and-freeze EM acute slice samples from a past study^23^. We acquired images of neurons synapsing directly onto dendritic shafts or neuronal cell bodies (somas). These synapses also displayed fainter postsynaptic densities and as such served as approximations of inhibitory synapses based on previously published EM definitions^30,38–41^. Notably, uncoated pits formed 100 ms after a single stimulus in these synapses (Figure 4A,B; see Figure S4 for more EM images and examples of approximated inhibitory synapses located on somas or projections). These pits formed primarily 8-46 nm from the active zone (Figure 4C, median pit distances: 46.23, 11.58, and 8.25 nm for mouse 1, 2, and 3 respectively). Clathrin-coated pits were occasionally present, but did not significantly increase in numbers after stimulation at the timepoints examined (Figure 4B). No apparent Ω-structures indicative of potential kiss-and-run like events^12,43,44^ were observed at synapses at any of the tested time points. Take together these results suggest a model where cortical inhibitory presynapses perform clathrin-independent endocytosis on a millisecond timescale following a single stimulus

**Figure 4.**
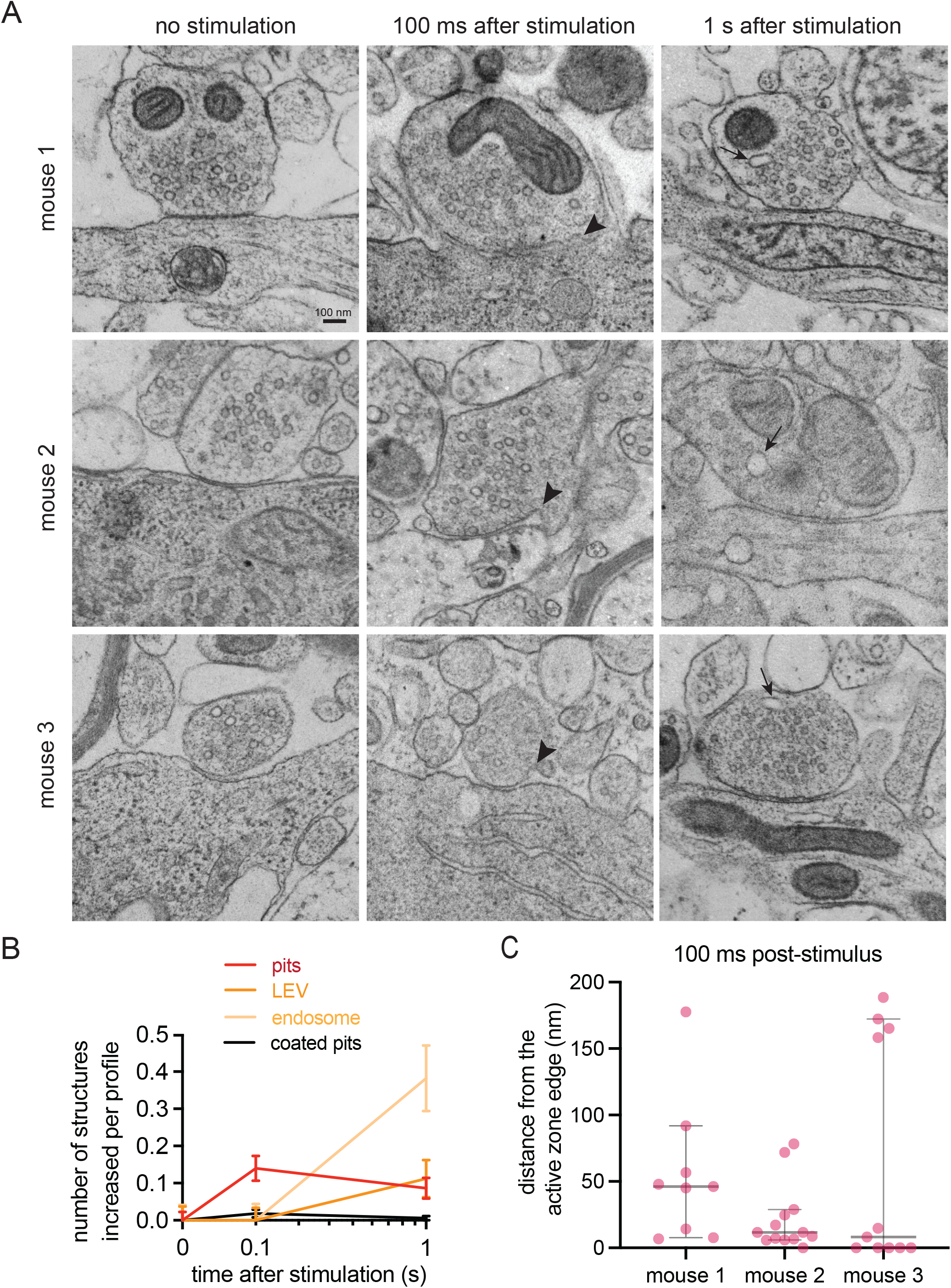
Uncoated pits form 100 ms after stimulation in putative inhibitory synapses. (A) Example electron micrographs showing uncoated endocytic pits (black arrowheads) and putative large endocytic vesicles and endosomes (black arrows) at the indicated time points in acute mouse cortical slices. Scale bar: 100 nm. More example TEM images are provided in Figure S4. (B) Plots showing the increase in number of each endocytic structure per synaptic profile after a single stimulus. Uncoated pits (pits), large vesicles (LEV), endosomes (endosome), and clathrin-coated pits (coated pits) are indicated. Data are pooled from slices from 3 independent mice (2 female and 1 male) and presented as mean ± SEM. n values for each timepoint: no stim, n = 150 synapses; 100 ms, n = 160 synapses; 1 s, n = 170 synapses. (C) Plot showing the distance distribution of uncoated endocytic pits from the edge of an active zone 100 ms post-stimulus in acute mouse cortical slices. Data are presented as median ± 95% confidence interval. Each dot represents an uncoated pit.

## Discussion

Using super-resolution and time-resolved EM imaging we explored the vesicle dynamics of inhibitory synapses. Similar to our recent findings in acute mouse and human cortical slices^23^, we find that cultured mouse cortical synapses form uncoated pits 100 milliseconds after a single stimulus. Increased ferritin positive large vesicle and endosomal structures confirm that endocytosis is occurring within milliseconds. Antibody labeling of endogenous Dyn1xA in cultured excitatory and inhibitory presynapses indicates that both subtypes contain a protein isoform necessary for ultrafast trafficking in the reported endocytic zone^25^. Specific examination of cortical inhibitory synapses in acute mouse brain slices recapitulates this Dyn1xA localization in a more intact tissue context. Cortical inhibitory synapses from acute mouse brain slices also exhibit uncoated pits forming 100 milliseconds post-stimulus. Our results suggest that inhibitory presynapses may exhibit ultrafast endocytic trafficking based on consistency with the definition of ultrafast endocytosis—uncoated pits forming 100 milliseconds after a single stimulation, within 100 nm from the active zone. These data seem to support a model where ultrafast endocytosis is a generalized pathway, though future experiments are warranted to further confirm if molecular requirements are similar.

As a preliminary proof of concept, we focused on ultrastructural dynamics in the timescale reported to initiate and complete ultrafast endocytic pit formation—100 milliseconds and 1 second. With a restricted timescale there are likely unseen differences in slower vesicle recycling mechanisms. Alongside existing reports illustrating that inhibitory synaptic vesicles reacidify slower^7,8^, our data may add that inhibitory vesicles are recycled fast within milliseconds, and reacidified at their own pace after uptake. Our data differ from studies performed near physiological temperature in hippocampal cultures, which indicate an approximate 80% slower endocytosis in inhibitory synapses^5^. These differences in results may be due to: our choice of testing cortical synapses as opposed to hippocampal, our use of single stimuli rather than repetitive stimuli, or our examination of membrane dynamics rather than pHluorin tagged proteins. Notably, some studies suggest about two-thirds of VGAT-labeled synaptic vesicles exhibit more prevalent kiss-and-run endocytosis behaviors^6^. It is interesting to theorize that to support extremely fast firing rates *in vivo*, like those found in fast-spiking cortical interneurons (some spiking upwards of 400 Hz in the mouse cortex^45^), synaptic transmission may be keeping up the pace through faster endocytosis mechanisms to maintain high-fidelity neural computations.

Our localization of endogenous Dyn1xA is in agreement with data illustrating that excitatory and inhibitory neurons express similar levels of Dynamin 1 protein^5^. Though we find Dyn1xA in both synapse types, there is no guarantee that the same protein is utilized similarly in excitatory and inhibitory synapses. An example of this phenomena has been seen with Synapsin induced clustering of synaptic vesicles—clustering vesicles via liquid-liquid phase separation in inhibitory synapses versus protein crosslinking in excitatory synapses^46^. Future studies must be done to examine the mechanistic details of Dyn1xA in inhibitory synapses. We also restricted our initial study to gephyrin-expressing inhibitory synapses. Gephyrin is a postsynaptic scaffold for synapses containing ionotropic, fast-acting GABA_A_ receptors^47^—though it is not necessary for GABA_A_ receptor clustering at all inhibitory synapses^48,49^. As such we did not investigate synapses containing slower, metabotropic GABA_B_ receptors^50^. Additionally, we do not distinguish the compositions of the gephyrin-expressing synapses we examined—as gephyrin is scaffolding for inhibitory glycine receptors^51^ as well, which are present in the mouse cortex^51,52^. Yet it is important to distinguish that glycinergic synapses can be both excitatory or inhibitory^53^, and potentially metabotropic^54^ alongside their canonical ionotropic state. It is crucial that future work take into account the complexity of inhibitory synapses to gain better insight into their physiological roles. Despite these limitations this brief study represents a step forward in dissecting the ultrastructural architecture of activated cortical microcircuits, excitatory and inhibitory.

## Acknowledgments

We would like to thank: Yuuta Imoto and Kie Itoh for the development of the Dyn1xA antibody and STED analysis code; Kie Itoh, Krishnaveni Jonnalagadda and Christian Pearson for preparation of neuron cultures; Sydney Brown for the assessment of preliminary EM images from cortical cultures; Barbara Smith, Mike Delanoy, LaToya Roker, Loza Lee, and Scot Kuo at the Johns Hopkins Microscopy Facility for technical and administrative assistance in electron microscopy; Aleksandr Smirnov and Susan McTeer at the Johns Hopkins Neuroscience Imaging Center for technical and administrative assistance in STED microscopy.

C.R.E. was supported by the Howard Hughes Medical Institute (HHMI) Gilliam Fellowship for Advanced Study (GT14961). S.W. was supported by funds from the Johns Hopkins University School of Medicine, Marine Biological Laboratory Whitman Fellowship, Brain Research Foundation Scientific Innovation Award, Helis Foundation award, the Kleberg Foundation grant, and the National Institutes of Health (R35 NS132153 and R01 MH139350) awarded to S.W. S.W. is an Alfred P. Sloan fellow, a McKnight Foundation Scholar, a Klingenstein and Simons Foundation scholar, and a Vallee Foundation Scholar. The EM ICE high-pressure freezer was purchased partly with funds from an equipment grant from the National Institutes of Health (S10RR026445) awarded to S. C. Kuo.

## Author contributions

### Chelsy Eddings

conceptualization, zap-and-freeze EM, TEM image acquisition and analysis, immunocytochemistry, immunohistochemistry, STED imaging and analysis, writing—original draft and figure construction

### Shigeki Watanabe

conceptualization, TEM image analysis, writing—original draft and figure construction

*all authors edited the manuscript

## Declaration of interests

The authors declare no competing interests.

## Figure legends

**Figure S1.** Additional STED images for Figure 1.

Overview 2D, three-color STED images of cultured mouse cortical synapses. Example side view (i and ii) and top view (iii and iv) synapses are highlighted as panels.

(A) Excitatory synapses observed on projections.

(B) Inhibitory synapses observed on a cell body (nucleus devoid of staining is denoted by * symbol).

(C Example Dyn1xA puncta in side view excitatory and inhibitory synapse images, which were present but not analyzed. Scale bars: 300 nm unless noted.

**Figure S2.** Additional EM images for Figure 2.

Example electron micrographs of cultured mouse cortical neurons that have undergone zap-and-freeze at the indicated time points. Uncoated endocytic pits (black arrowheads) and ferritin-positive membrane structures (black arrows) are indicated. Here no distinction was made between excitatory or inhibitory synapses. Scale bars: 100 nm.

**Figure S3.** Additional STED images for Figure 3.

(A) Overview 2D, three-color STED image of a cortical region in an acute mouse brain slice. Example side view (i and ii) and top view (iii and iv) synapses are highlighted as panels. Nuclei devoid of staining are denoted by * symbols.

(B) Example Dyn1xA puncta in side view inhibitory synapse images, which were present but not analyzed. Scale bar: 300 nm unless noted.

**Figure S4.** Additional EM images for Figure 4.

Example electron micrographs of acute mouse brain slices that have undergone zap-and-freeze. Synapses shown with non-prominent postsynaptic densities onto either (A) projection shafts or (B) somas/cell bodies.

(C) More example uncoated pits (black arrowheads) found 100 ms post-stimulus in synapses with non-prominent postsynaptic densities. Scale bars: 100 nm.

## Notes

### Competing Interest Statement

The authors have declared no competing interest.

