## Supplementary figures and images for "Ultrafast endocytosis in mouse cortical inhibitory synapses"

### supplemental figures

Figure S1. Eddings, et al.

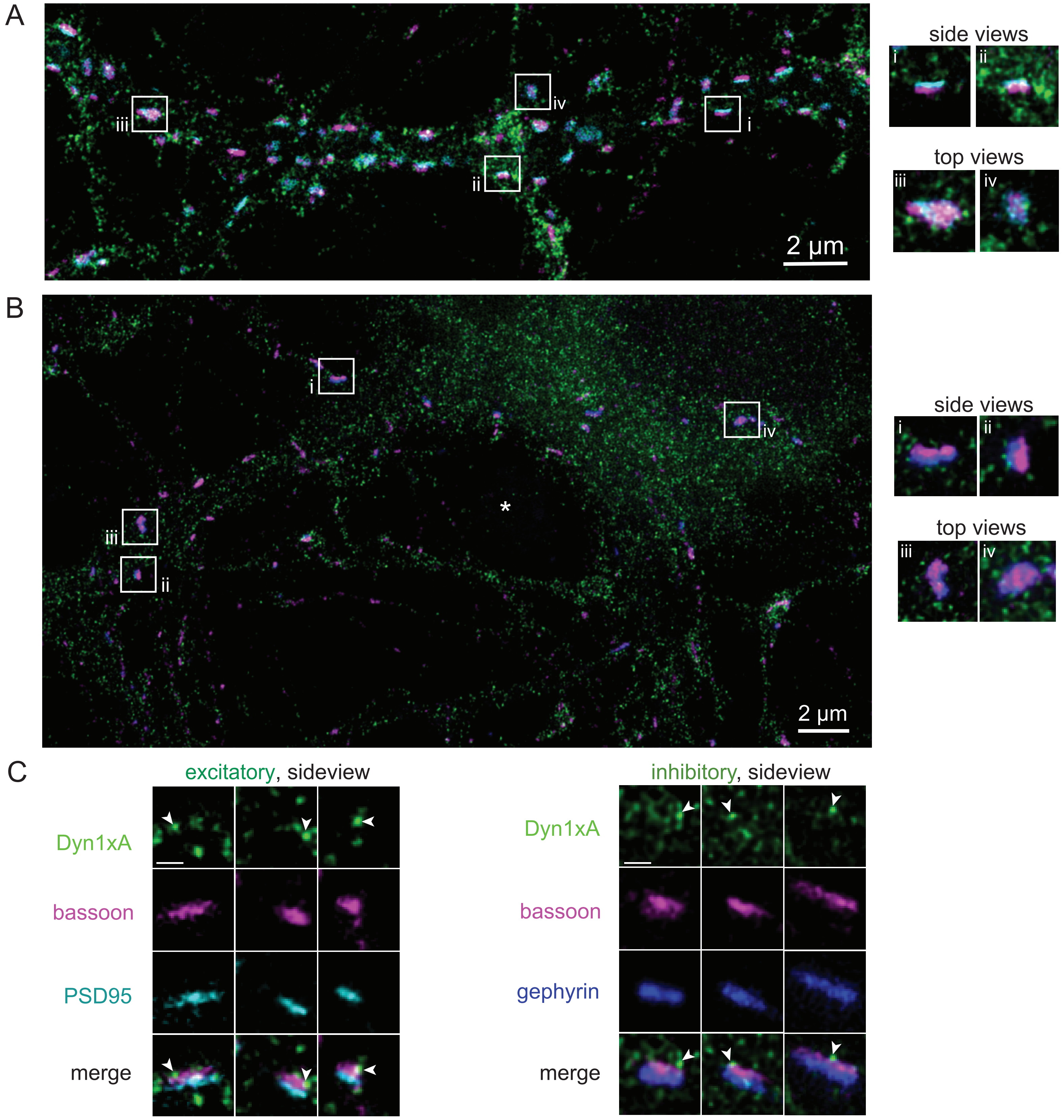

Figure S2. Eddings, et al.

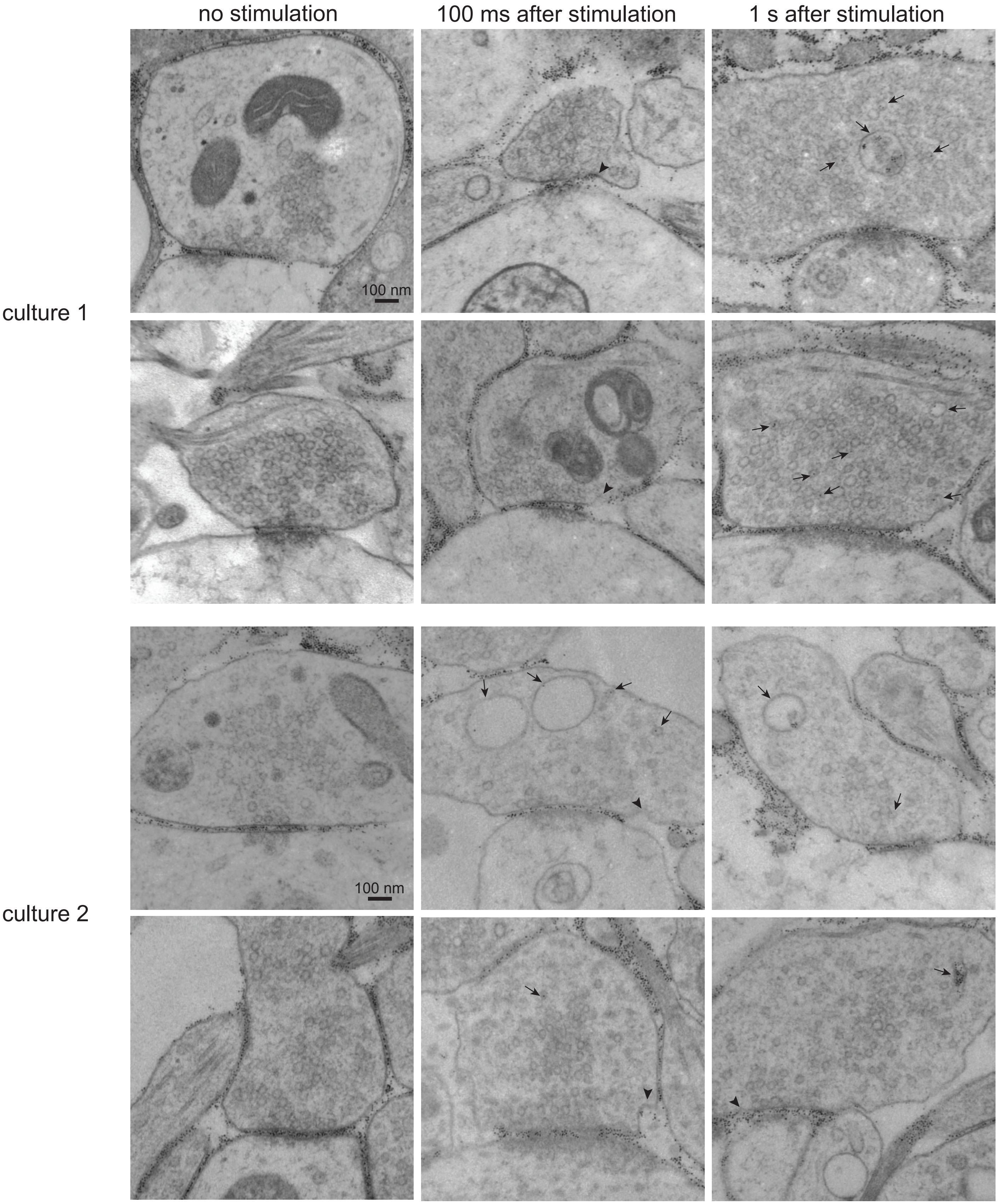

Figure S3. Eddings, et al.

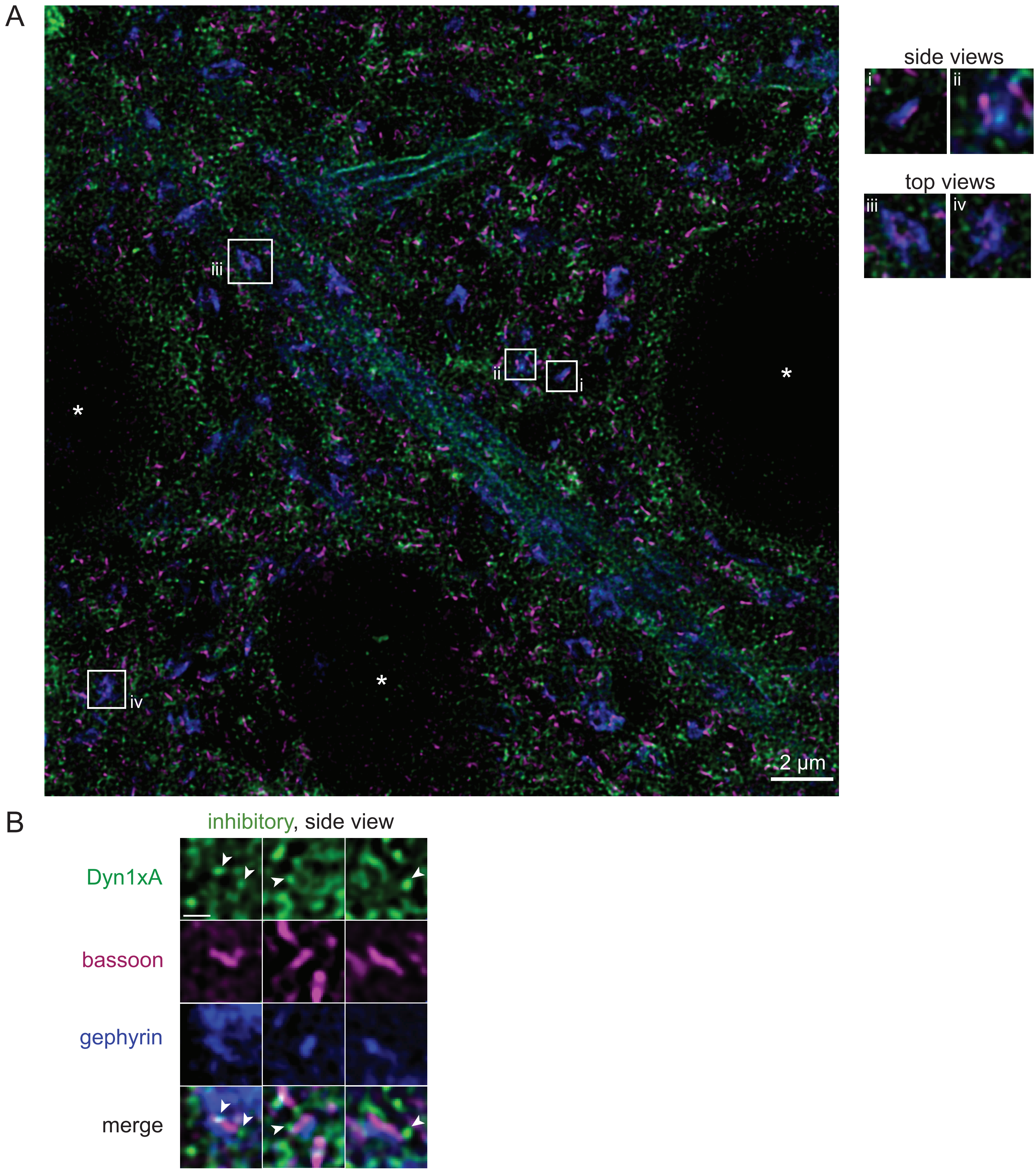

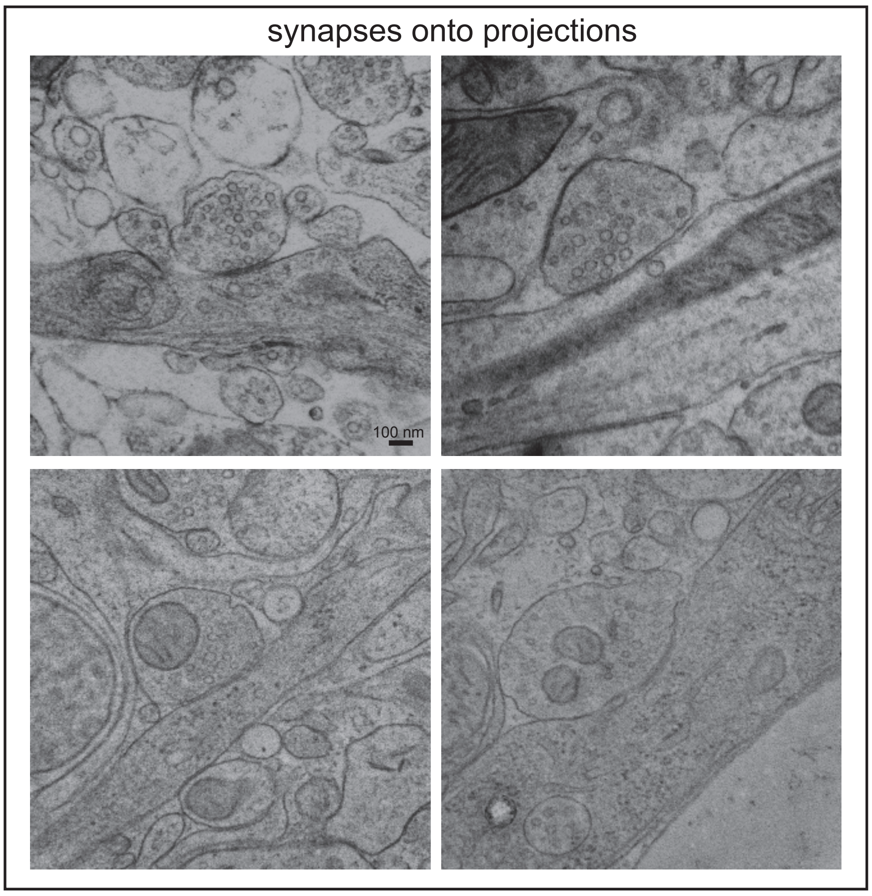

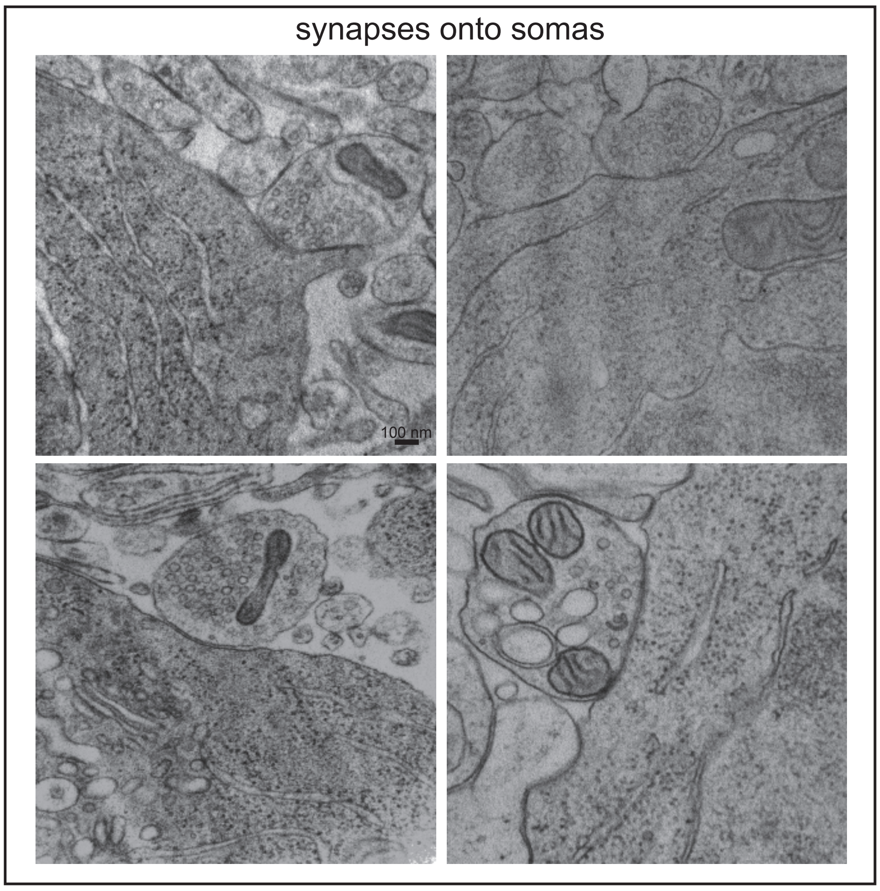

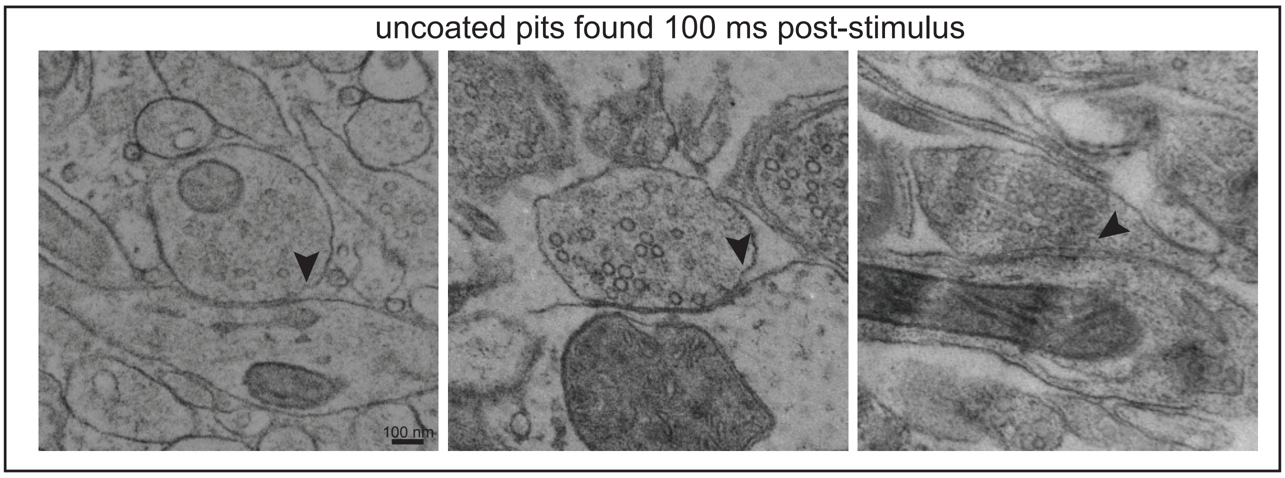
